# Incorporation of plant residue in corn monoculture selectively enriches antagonistic phenotypes among lignin-degrading vs non-degrading *Streptomyces*

**DOI:** 10.1101/2025.07.14.664736

**Authors:** Brett R Lane, Angela K Tran, Miriam F Gieske, Linda L Kinkel

**Affiliations:** Department of Agronomy, Kansas State University, Manhattan, Kansas, USA; Department of Plant Pathology, University of Minnesota, St. Paul, Minnesota, USA; Department of Biology, University of Minnesota Morris, Morris, Minnesota, USA

**Keywords:** *Streptomyces*, Resource competition, Antagonism, Lignin

## Abstract

Competition for limited carbon resources in the soil can drive complex microbe-microbe dynamics including selection for antagonistic resource competitors. Lignin, one of the main components of plant residue, is one source of carbon input into soil. However, the highly recalcitrant nature of lignin requires degrading organisms to invest in extracellular enzymes, whose breakdown products are susceptible to poaching by opportunistic competitors. Here, we tested two hypotheses: 1) lignin-degrading *Streptomyces* are more inhibitory than non-degrading isolates, and 2) competition for lignin contributes to the frequency and intensity of antagonistic phenotypes. We collected 134 *Streptomyces* from the soil of 55-year corn monocultures that had been managed in a +/− fertilizer by +/− residue removal factorial design. Using a subset of 24 lignin-utilizing and 24 non-utilizing isolates we measured the inhibition of each isolate against ten *Streptomyces* standards. Lignin-degrading *Streptomyces* had larger mean zones of inhibition than non-lignin-degrading populations, but only in plots where residue was retained, suggesting competition for lignin promotes the evolution or maintenance of antagonistic interactions when plant residues provide abundant lignin substrates. Among non-lignin-degrading *Streptomyces*, there was no difference in the frequency or intensity of antagonistic phenotypes with fertilizer or residue practices. Our results suggest that competition for byproducts of lignin degradation by lignin-degrading *Streptomyces* imposes selection for antagonistic phenotypes in habitats with regular residue inputs, but not in non-lignin-enriched soils. This work has potential implications for the development of antibiotic-mediated disease suppressive soils, the survival of plant pathogens within residue, and the structure and function of the soil microbial community.

## Introduction

The production of antimicrobial secondary metabolites by soil microbes is one of the mechanisms that can contribute to the development of naturally disease-suppressive soils. The role of secondary metabolites in disease suppression has been well-studied (Kinkel et al. 2011; Lan et al. 2024; Li et al. 2024; Kinkel et al. 2012, 2014). Understanding the mechanisms that drive selection for the capacity of soil-borne bacteria, including *Streptomyces*, to produce antimicrobial secondary metabolites is critical to promoting the development of sustainable antimicrobial secondary metabolite-mediated disease suppressive soils.

The association of disease suppressive soils with long-term monocultures (Mazzola 2002; Schlatter et al. 2017; Weller et al. 2002; De La Fuente et al. 2006) has prompted the hypothesis that low diversity of root exudates of crop monocultures may stimulate selection for antagonistic phenotypes as a consequence of competition for a small number of carbon resources (Kinkel et al. 2011). Support for this hypothesis was observed in work showing enrichment of antagonistic *Streptomyces* populations following amendment of single carbon substrates, but not amendment with water or multiple carbon substrates (Schlatter et al. 2009; Dundore-Arias et al. 2019).

Further, the addition of glucose, fructose, malic acid, or a mixture of these substrates to low organic matter soil mesocosms increased the frequency and intensity of *Streptomyces* inhibitory phenotypes against plant pathogenic organisms, supporting the hypothesis that resource competition may facilitate the development of disease suppressive soils (Dundore-Arias et al. 2020).

Lignin, which accounts for 30% of all organic carbon in the biosphere (Ez-Zahraoui et al. 2023), is a highly recalcitrant organic polymer that is primarily incorporated into agronomic soils through crop residues (Rasse et al. 2006). Despite being regarded as one of the most stable components of soil organic matter, a growing body of evidence has established that 90% of lignin within the soil is metabolized rather than being incorporated into long-term stable, soil carbon (Rasse et al. 2006; Hofmann et al. 2009; Dignac et al. 2005; Heim and Schmidt 2007; Kiem and Kögel-Knabner 2003). In forest settings, lignin is degraded primarily by white-rot fungi, the only group capable of completely mineralizing lignin (Shah et al. 1992). In comparison, relatively little research has explored the degradation of lignin in agronomic settings. Interestingly, the soil lignin content of arable soil is one-and-a-half times that of grassland soils and nearly twice that of forest soils, possibly as the conditions in arable soils (pH and the inorganic nitrogen fertilizers) are less conducive for lignin-degrading white-rot fungi than forest soils (Pradeep Kumar et al. 2023; Thevenot et al. 2010). Yet the turnover of lignin within arable soils indicates that lignin in these soils is degraded, whether by white-rot fungi or by consortia of other microorganisms including fungi and bacteria (Thevenot et al. 2010).

One of the most prominent lignin-degrading bacterial groups is the actinomycetes (Brown and Chang 2014), including the genus *Streptomyces*. *Streptomyces* are filamentous, soil-borne, Gram-positive bacteria that are known for their extensive production of antimicrobial secondary metabolites. While the metabolic pathways of lignin-degrading bacteria are not as well-studied as for fungi, work on the model lignin-degrading bacteria *Streptomyces viridosporus* T7A suggests the bacterial degradation of lignin relies on the secretion of extracellular peroxidases to cleave aromatics from the lignin polymer (Thomas and Crawford 1998; Ramachandra et al. 1988). Once secreted, these extracellular enzymes may function as a public good, with their products open to consumption by co-occurring, non-lignin-degrading organisms (‘cheaters’). In the absence of measures to protect these products, enzyme producers, hampered by the cost of enzyme production, may face competitive exclusion by cheaters (Allison 2005). Thus, lignin-degrading microbes, such as *Streptomyces*, may face selection to protect their investment in extracellular enzymes (Schimel and Weintraub 2003; Allison et al. 2010).

Variation in lignin inputs and turnover across agricultural management systems provides a tractable axis along which these selective pressures may operate. In particular, residue incorporation directly alters the quantity and spatial distribution of lignocellulosic substrates, while fertilization can influence carbon availability in soils as well as microbial demand for carbon and the activity of lignin-degrading communities (Rasse et al. 2006). These factors together are expected to regulate not only the abundance of lignin-derived compounds, but also the extent to which their products are contested resources within the soil matrix. Under conditions where lignin metabolism generates localized, competition-prone pools of aromatics, selection may favor organisms capable of both accessing these substrates and defending them, whereas reduced lignin inputs or more diffuse resource environments may weaken such selection. Thus, management-driven differences in lignin dynamics provide a mechanistic basis to evaluate how resource complexity and accessibility shape the prevalence of antagonistic phenotypes in lignin-utilizing microbial populations.

Here, we tested two hypotheses: 1) *Streptomyces* that utilize lignin would be significantly more inhibitory than *Streptomyces* that cannot utilize lignin, and 2) competition for locally-derived lignin incorporated into the soil influences the frequency and intensity of antagonistic *Streptomyces* phenotypes. In this work, we studied soil *Streptomyces* within long-term (55 year) maize monocultures with or without annual fertilizer applications factorial with or without residue incorporation. This work sheds light on the impacts of lignocellulosic materials as mediated by cultural management practices on the functions of microbial populations in agronomic soils.

## Methods

### Experimental design and soil sampling

We sampled soils from maize plots at the Rosemount Research and Outreach Center that have been maintained in continuous monoculture since 1959. In these plots fertilizer application and crop residue retention/removal have been manipulated in a factorial manner for the lifespan of the continuous monoculture. Each plot was replicated for a total of two blocks.

Soil samples were collected in November 2015 as described by Gieske and Kinkel (Gieske and Kinkel 2020). In brief, 15 m x 15 m plots were divided into four 7.5 m x 7.5 m quadrants. Within each quadrant, a single soil sample (comprised of 3 bulked soil cores, each 2 cm diameter x 15 cm depth) was collected from a random location within the quadrant. Samples from all plots were transported to the lab on ice and stored for up to 4 days at 4°C before processing.

### Streptomyces isolates

*Streptomyces* strains were isolated from soil as described by Gieske and Kinkel (Gieske and Kinkel 2020). In brief, we air-dried soil samples at room temperature under sterile cheesecloth and passed each sample through a 2 mm sieve to remove plant material. We then placed five grams of each dried sample in 50 mL of sterile deionized water and shook (175 rotations per minute) each sample for one hour at 4°C. Soil suspensions were serially diluted with sterile deionized water to 10^−2^ and 50 uL of each dilution was spread onto three replicate starch casein agar plates (Gieske and Kinkel 2020). After 6-9 days incubation at 28°C, we transferred five random colonies (per soil sample) with characteristic *Streptomyces* morphology to Oatmeal Agar (OA). Isolates were grown at 28°C for 3-7 days and subcultured as necessary to ensure purity. Purified isolates were stored at −20°C in 20% glycerol.

### Resource use characterization

Nutrient use phenotypes of each Streptomyces isolate were determined using Biolog SF-P2 plates (Biolog Inc.) as described by Gieske (Gieske 2019) approach characterizes isolate growth on 95 distinct carbon sources by measuring the optical density (OD) at 590 nm in relation to a water control. In brief, we grew *Streptomyces* isolates on OA plates at 28°C for 7 days or until thick spores were observed. We collected spores into 0.2% carrageenan using a sterile cotton swab. We adjusted suspensions to an OD (590 nm) of 0.20-0.24 before diluting 1.5 mL of the spore suspension into 13.5 mL of sterilized 0.2% carrageenan for a total of 15 mL. We inoculated Biolog SF-P2 plates with 100 µL per well of spore suspension. The capacity of an isolate to utilize each carbon source was determined by measuring the absorbance of each well at 590 nm after 72 h of growth at 28°C. The absorbance of the control well (no carbon substrate) was subtracted from each other well. Wells with a computed OD of less than 0.005 were transformed to 0 (Gieske 2019).

To determine the ability of *Streptomyces* to utilize lignin as a sole carbon source we tested the ability of isolates to grow on lignin minimal media (lignin MM). Minimal media was adapted from ActinoBase.org and previous work (Hopwood 1967; Lane et al. 2023) and consisted of (per liter deionized water) 5 g ammonium sulfate, 2 g potassium nitrate, 2 g sodium chloride, 2 g dipotassium phosphate, 0.05 g magnesium sulfate heptahydrate, 0.01 g ferrous sulfate heptahydrate, 0.001 g zinc chloride, 15 g of agarose, and 5 g of the selected carbon source (kraft lignin or glucose). Agarose was used in place of agar as some *Streptomyces* can grow on agar as a sole carbon source. In brief, we dotted 10 µL of freezer stocks on lignin minimal media and allowed the dot to dry. After 7 days of growth at 28°C a 5 mm x 5 mm agarose plug, containing none of the original dotted material, was cut from the colony tip and placed face down on a subsequent plate of lignin MM. *Streptomyces* that had expanded outward from the agar plug and into the second generation of lignin MM after 7 days growth were determined to be able to utilize lignin as a sole carbon source. Isolates were concurrently plated on glucose MM and MM absent any additional carbon source to prevent false negatives and positives.

### Characterization of inhibition and resistance phenotypes

We selected three lignin-utilizing and three non-utilizing Streptomyces isolates from each plot continuously cropped to maize for characterization of inhibitory and resistance phenotypes. These forty-eight focal *Streptomyces* isolates were selected to represent the full factorial design of the experiment (3 isolates per plot x 2 fertilization practices x 2 residue treatments x 2 blocks x the (in)ability to utilize lignin as a sole carbon source) while otherwise being selected at random.

We determined the inhibitory phenotype of each focal *Streptomyces* isolate using a modification of the agar overlay method described by Kinkel et al. (2014). In brief, we dotted 4 µL of a spore suspension onto 15 mL SCA plates and incubated at 28°C for 3 days. Dotted isolates were killed by inverting the petri plates over a watch glass containing 4 mL of chloroform for 1 h. Plates were then uncovered in a laminar flow hood for 30 minutes to allow the evaporation of residual chloroform. We then overlaid each plate with 10 mL of cooled, molten SCA before spreading 100 µL of one of ten target *Streptomyces* isolates on the solidified and cooled layer. Plates were incubated at 28°C for 3 days. We measured the size of any zone of inhibition in mm from the edge of the dotted colony to the edge of the cleared zone. Any zone of inhibition for an individual isolate dot was measured twice, at perpendicular angles, and averaged. Each focal-target isolate pair was replicated three times. The intensity of inhibition for each isolate pair was defined as the average zone of inhibition across all replicates. Only zones of inhibition averaging greater than 2 mm across all replicates were considered to be inhibitory.

We characterized the resistance of focal *Streptomyces* to common clinical antibiotics using disc diffusion assays - method described by Schlatter and Kinkel (D. C. Schlatter and Kinkel 2014). In brief, 100 µL of a focal isolate spore suspension was spread onto 15 mL SCA plates and allowed to dry before filter paper discs containing a known quantity of an antibiotic (Table S1) (BD BBL Sensi-Disc; Becton, Dickinson and Company, Sparks, MD) were placed onto the dried isolate overlay. Inhibition zones were measured as described above after 3 days of growth at 28°C. Inhibition zones averaging less than 2 mm were considered fully resistant.

### Isolate sequencing and phylogenetic analysis

The 16S rRNA gene was previously sequenced and assigned taxonomy via manual inspect of BLAST by Gieske and Kinkel (2020). Here, we aligned sequences (available as a fasta in the supplement) using the default settings of the *Geneious Alignment* function within the Geneious Prime program [v2025.1.2] with automatic determination of read direction enabled.

Alignments were manually inspected for quality. A single non-*Streptomyces* isolate, identified as a *Nocardia* species, was included in the alignment to serve as an outgroup. To form a maximum-likelihood phylogenetic tree in the R package *phangorn* [version 2.12.1] (Schliep 2011) we used the function *modelTest* to select the TIM2 with gamma distributions and invariable sites (Posada 2008). We then formed the maximum-likelihood tree by feeding the output of *modelTest* to the function *pml_bb*. We then removed the outgroup and tested the phylogenetic clustering of *Streptomyces* isolates by management practice (residue and fertilizer treatments), block, and the ability to utilize lignin as a sole carbon source using Pagel’s lambda statistic and 10,000 simulations as implemented in the *phylosig* function in the R package *phytools* [v2.4.4] (Pagel 1999; Revell 2024). Zero-length terminal branches (exact matches within the alignment) were transformed by adding one hundredth of a percent of the smallest non-zero terminal branch length; thus, Pagel’s lambda was selected over Bloomberg’s K due to the known susceptibility of Bloomberg’s K (and the robustness of Pagel’s lambda) to type I biases when faced with incompletely resolved phylogenies and suboptimal branch-length information (Molina-Venegas and Rodríguez 2017). Visualizations of the phylogenetic tree were made using the packages ggplot2 [v3.5.1] and ggtree [v3.14.0] (Yu et al. 2017; Wickham 2016).

### Statistical analysis

All statistical analyses were performed in R version 4.4.2 using the package *lme4* [v1.1.37], with p-values provided by the package *lmerTest* [v3.1.3] (Bates et al. 2015; Kuznetsova et al. 2017). The normality of residuals was confirmed through inspection of residual plots. We tested the impact of fertilizer addition and residue removal/retention on the ability of 134 isolates to grow on lignin as a sole carbon source using a binomial generalized linear regression using block as a random intercept.

We tested the impact of the ability to utilize lignin as a sole carbon source and cultural management practices on the nutrient use profiles of *Streptomyces* using a PERMANOVA with 10,000 permutations, stratified by block, as implemented by the function *adonis2* in the R package *vegan* [v2.6.10] (Oksanen et al. 2022). The distance matrix was calculated using Euclidean distances. The niche width was determined as the number of utilized carbon sources (OD>0.005) after 72 hours. We calculated the average growth of each isolate as the average OD of all utilized carbon sources after 72 hours. We also tested the impact of the ability to utilize lignin as a sole carbon source and cultural management practices on the niche width and average growth using a linear regression with block as a random intercept. We determined if the ability to utilize lignin as a sole carbon source was associated with changes in the growth of isolates on each of the 95 carbon sources individually (when OD>0.005) using a linear regression with a Bonferroni multiple comparison correction.

The intensity of inhibition for a given focal *Streptomyces* isolate was calculated as the average zone of inhibition across all standards that were inhibited (>2 mm zone of inhibition) (Dundore-Arias et al. 2019; Schlatter and Kinkel 2014). We determined the impact of the ability of focal isolates to utilize lignin and cultural management practices on the inverse square root transformed intensity of inhibition using a linear regression with block as a random intercept.

Visualizations of the intensity of inhibition were conducted using a log10 transformation rather than the inverse square root for ease of interpretation. Cohen’s D for significant pairwise comparisons was calculated using the function *cohensD* in the R package *lsr* [v0.5.2] (Navarro 2015). We also tested the impact of cultural management practices on the intensity of inhibition within lignin utilizing and non-utilizing *Streptomyces* using a robust linear regression as implemented by the *rlm* function within the R package *MASS* [v7.3.65] with p-values provided by the *rob.pvals* function within the *repmod* R package [v0.1.7] (Venables et al. 2002). The impact of the ability of focal isolates to utilize lignin and cultural management practices on the proportion of standards inhibited was tested using a binomial generalized linear model with block as a random intercept.

We tested the impact of the ability of focal isolates to utilize lignin and cultural management practices on the proportion of isolates that were inhibited by antibiotic discs (>2 mm zone of inhibition translated into binary inhibit/not inhibited) using a binomial generalized linear regression with block as a random intercept. We also tested the ability of focal isolates to utilize lignin and cultural management practices on the square root transformed zone of inhibition for isolate-antibiotic combinations that were not completely resisted (inhibition zones > 2 mm). These comparisons were conducted using a linear regression with antibiotic identity as a random intercept. Antibiotic identity was used in place of block as a random intercept as the concentration of each antibiotic differed.

## Results

### Lignin use phenotypes are phylogenetically clustered within Streptomyces

Forty-four of 134 *Streptomyces* isolates (32.8%) were able to utilize lignin as a sole carbon source on *Streptomyces* minimal medium. The proportion of *Streptomyces* that could use lignin as a sole carbon source was significantly greater in *Streptomyces* isolated from plots that had received fertilizer (37.9% utilize lignin) than plots that did not receive fertilizer (27.9%) (Table 1) but was not impacted by residue practices. Phylogenetic analysis of the 16S region indicated that the ability to utilize lignin as a sole carbon source was phylogenetically clustered (Figure 1). Pagel’s lambda confirmed the phylogenetic clustering of lignin use (Table S2). The fertilization practices of plots from which isolates were collected was also reflected in the phylogenetic clustering of isolates (Table S2). Isolates were not phylogenetically clustered by residue treatment or block (Table S2).

**Figure 1.**
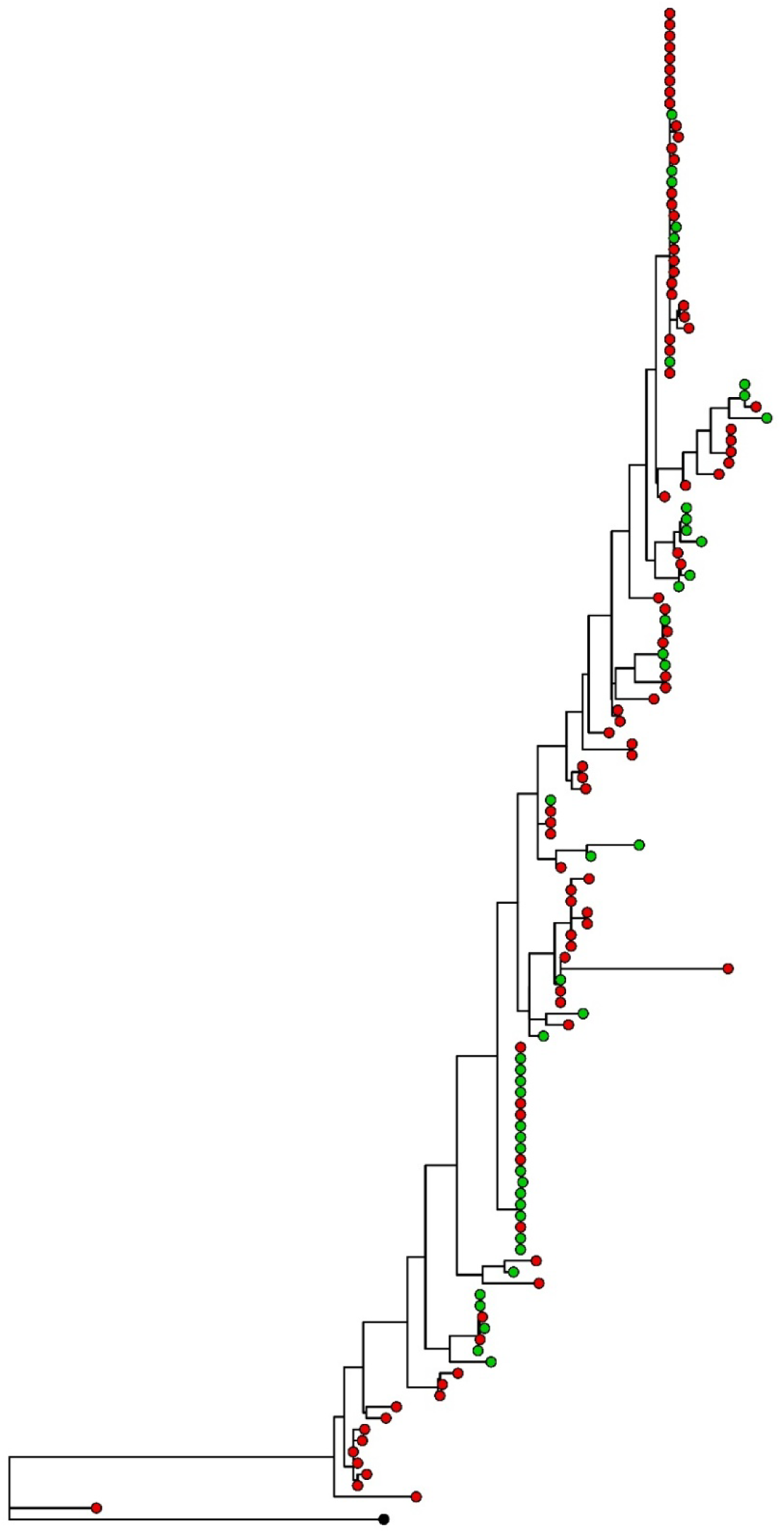
Maximum likelihood phylogenetic tree formed from the 16S region of *Streptomyces* isolated obtained from long-term corn monocultures. Green tips indicate the ability to utilize lignin as a sole carbon source while red tips indicate the isolate is unable to utilize lignin. The single isolate with a black tip signifies the outgroup (*Nocardia sp*.), which was not tested for lignin utilization.

**Table 1.** Binomial generalized linear model of the impact of fertilizer, residue, and their interaction on the ability of isolates to utilize lignin as a sole carbon source. The block from which isolates were obtained was used as a random effect. 37.9% and 27.9% of *Streptomyces* isolates from plots which did and did not receive fertilizer applications, respectively, were able to utilize lignin as a sole carbon source. Bold values indicate significance.

|  | Z-value | P-value |
| --- | --- | --- |
| Fertilizer | 2.025 | **0.0429** |
| Residue | 0.184 | 0.854 |
| Fertilizer x Residue | -1.540 | 0.124 |

### Lignin-utilizing *Streptomyces* exhibit greater intensity of antagonism

We found that the intensity of inhibition (average zone of inhibition by an isolate against allopatric isolates) by *Streptomyces* depended upon both whether the isolate was able to utilize lignin as a sole carbon source and whether the isolate came from residue-amended versus non-amended soil (Table 2, Figure 2). Specifically, lignin-utilizing and non-utilizing *Streptomyces* did not differ in the intensity of inhibition in plots when originating from plots that had their residue removed at the end of each growing season (Table S3). However, when residue was retained, *Streptomyces* that could utilize lignin as a sole carbon source were significantly more inhibitory than isolates that could not (Table S3). Interestingly, fertilizer application weakened the strength of the association between lignin utilization and the strength of inhibition (Cohen’s D fertilizer– = 1.602, Cohen’s D fertilizer+ = 1.020; Table S3; Figure 2). Residue practices as well as the interaction of fertilizer and residue practices, were associated with significant changes in the intensity of inhibition for lignin-utilizing *Streptomyces* (Table S4; Figure 2). Cultural management practices did not impact the intensity of inhibition among *Streptomyces* that cannot utilize lignin as a sole carbon source (Table S4; Figure 2).

**Figure 2.**
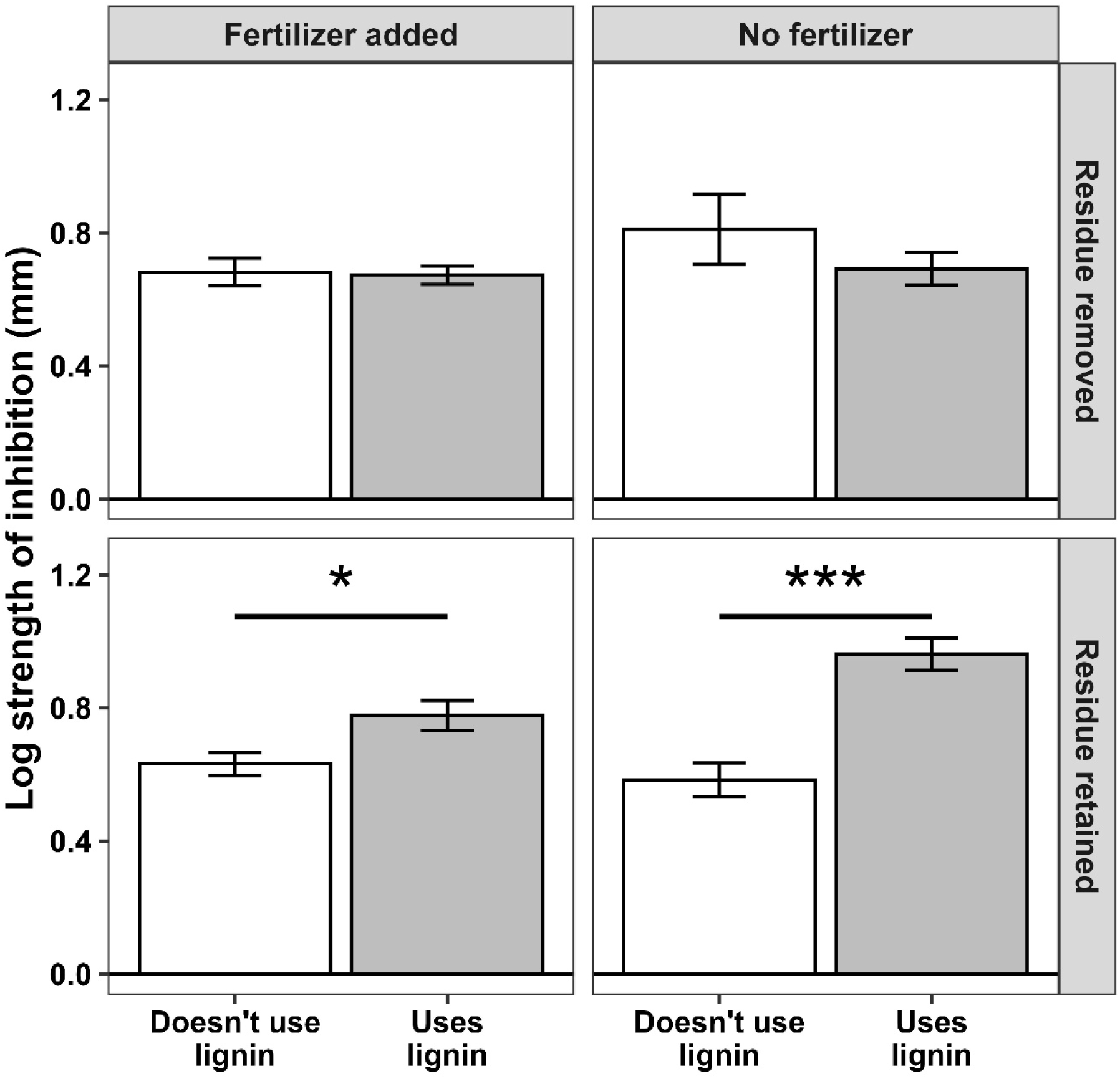
Log strength of inhibition (measured in mm of inhibition) of lignin utilizing and non-utilizing *Streptomyces*. Bars indicate standard error. Stars indicate significance: *p<0.05, **p<0.01, ***p<0.001.

**Table 2.** Binomial generalized linear model (left) and linear model (right) of the impact of fertilizer application, residue removal/retention, and the ability of *Streptomyces* to utilize lignin as a sole carbon source on the number of standards a given focal *Streptomyces* isolate could inhibit (left) and the inverse square root intensity of inhibition in inhibitory interactions (right). The block from which isolates were obtained was used as a random effect. Bold values indicate significance.

|  | Number of standards inhibited |  | Intensity of inhibition (zone size) |  |
| --- | --- | --- | --- | --- |
|  | Z-value | P-value | F-value | P-value |
| Lignin† | 0.625 | 0.532 | 8.664 | **0.00390** |
| Fertilizer | 0 | 1 | 0.0756 | 0.784 |
| Residue | 0 | 1 | 4.521 | **0.0355** |
| Lignin x Fertilizer | 0 | 1 | 1.359 | 0.246 |
| Lignin x Residue | 1.603 | 0.109 | 13.634 | **0.000337** |
| Fertilizer x Residue | -0.487 | 0.626 | 0.948 | 0.332 |
| Lignin x Fertilizer x Residue | -1.156 | 0.248 | 2.774 | 0.0984 |

†The ability of a *Streptomyces* isolate to utilize lignin as a sole carbon source

In contrast with the intensity of inhibition, the frequency of inhibition did not vary with management. The number of standards inhibited by individual *Streptomyces* ranged from 0 to 8 (of 10). Eight isolates (of 48) inhibited 7 or more standards while 38 isolates inhibited three or fewer standards. Our binomial generalized linear model indicated the number of standards inhibited was not associated with cultural management practices, the ability of an isolate to utilize lignin, or their interactions (Table 2).

### Lignin-utilizing *Streptomyces* resist inhibition by antibiotics

*Streptomyces* isolates exhibited complete resistance [<2 mm inhibition] to from 0 to 3 (of 10) antibiotics, with 43 of 48 isolates resisting 0 or 1 antibiotics. Neither the ability to utilize lignin nor cultural management practices were associated with the number of antibiotics resisted (Table 3). We also tested the size of the zone of inhibition of isolates that exhibited incomplete resistance to an antibiotic. *Streptomyces* that utilize lignin as a sole carbon source were significantly more resistant to antibiotics (smaller inhibition zone sizes) than *Streptomyces* that could not utilize lignin (Table 3). Post-hoc analysis revealed that while lignin-utilizing *Streptomyces* exhibited greater resistance (smaller zone of inhibition) against all ten antibiotics tested, differences were significant for only two of the antibiotics tested (chloramphenicol and rifampin; Figure S1).

**Table 3.** Binomial generalized linear model (left) and linear model (right) of the impact of fertilizer application, residue removal/retention, and the ability of *Streptomyces* to utilize lignin as a sole carbon source on the number of antibiotics a given focal *Streptomyces* isolate could completely resist (left) and the square root intensity of inhibition by antibiotics in interactions which did not exhibit complete resistance (right). The antibiotic used in each interaction was used as a random effect. Bold values indicate significance.

|  | Complete resistance |  | Zone size of resistance |  |
| --- | --- | --- | --- | --- |
|  | Z-value | P-value | F-value | P-value |
| Lignin† | 0 | 1 | 12.633 | **0.000421** |
| Fertilizer | 1.396 | 0.163 | 0.477 | 0.490 |
| Residue | 0.454 | 0.650 | 0.0655 | 0.798 |
| Lignin x Fertilizer | 0 | 1 | 1.164 | 0.281 |
| Lignin x Residue | 0.238 | 0.812 | 0.0622 | 0.803 |
| Fertilizer x Residue | -0.752 | 0.452 | 2.516 | 0.113 |
| Lignin x Fertilizer x Residue | 0.0870 | 0.931 | 1.131 | 0.288 |

†The ability of a *Streptomyces* isolate to utilize lignin as a sole carbon source

### Lignin utilization is associated with shifts in *Streptomyces* nutrient use

Lignin use, but not cultural management practices, had a significant impact on *Streptomyces* nutrient use profiles after 72 hours of growth on Biolog-SF-P2 plates (Figure 3, Table 4). We also investigated the impact of lignin utilization and cultural management practices on niche width (number of Biolog SF-P2 carbon sources with an OD>0.005) and average growth (average OD across utilized [OD>0.005] carbon sources) of *Streptomyces*. After 72 hours of growth, *Streptomyces* isolates that could utilize lignin exhibited significantly more growth than isolates that could not utilize lignin (Table 4). Post-hoc analysis with multiple comparison corrections revealed shifts in the average growth of *Streptomyces* could be attributed to D-gluconic acid, on which *Streptomyces* that could use lignin grew to a higher OD (Figure S2).

**Figure 3.**
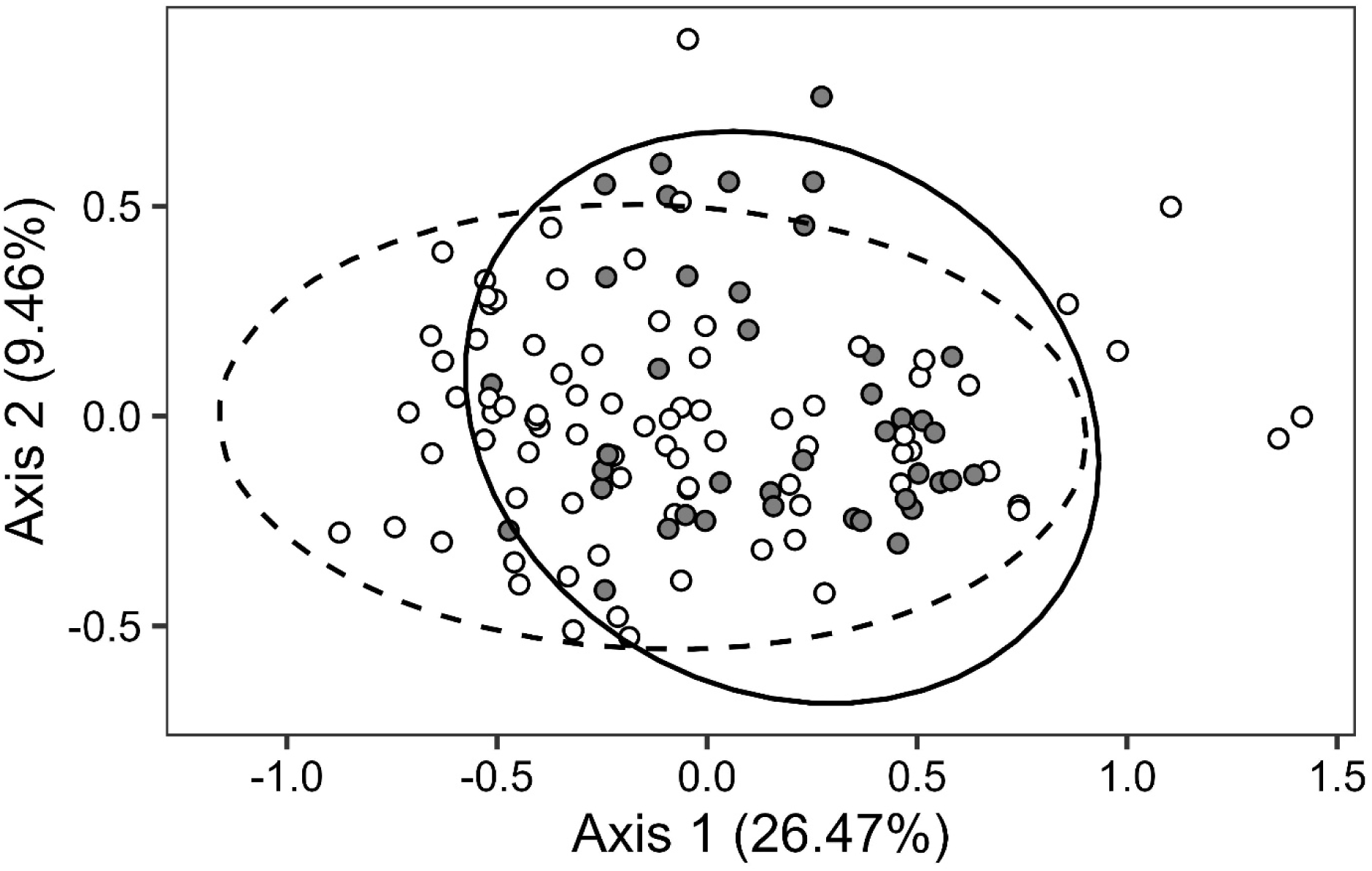
Principal Coordinates Analysis of *Streptomyces* nutrient use profiles as measured by Biolog SF-P2 plates (95 carbon sources) after 72 hours of growth. Grey dots and the solid line are *Streptomyces* which can utilize lignin while white dots with the dashed line cannot utilize lignin.

**Table 4.**
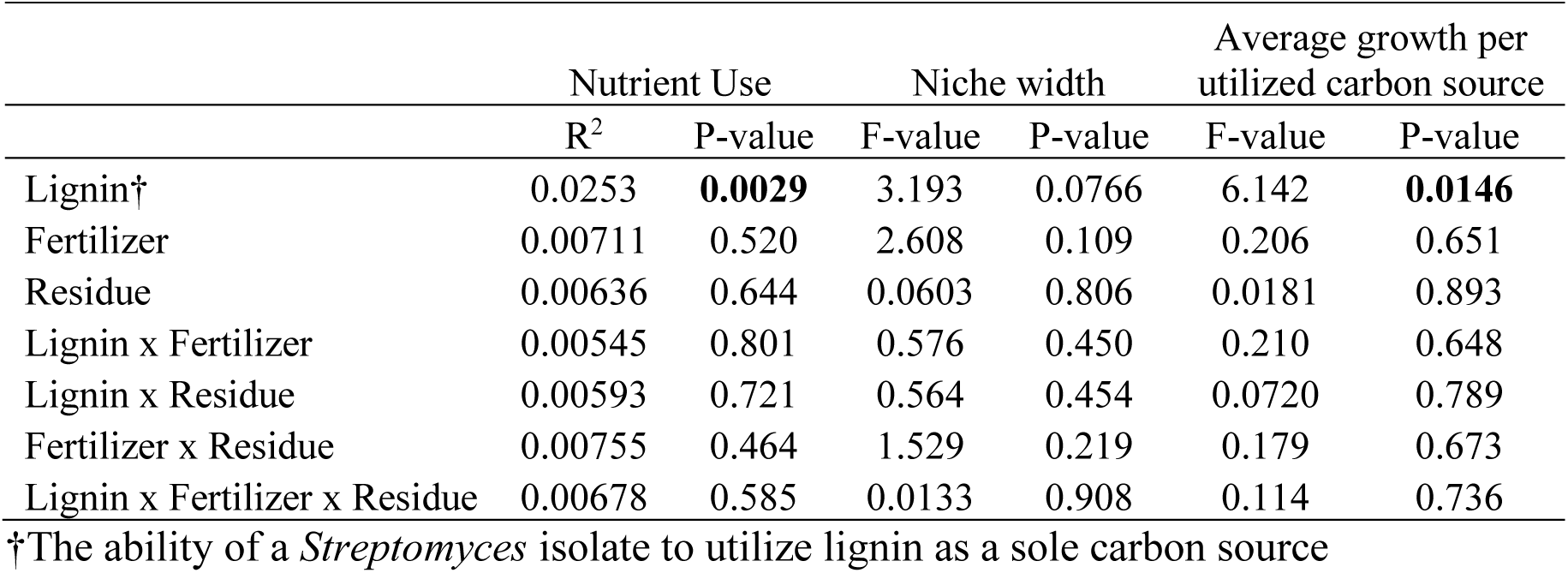
Impact of the ability of *Streptomyces* isolates to utilize lignin as a sole carbon source, residue removal/retention, and fertilizer application on the nutrient use profiles, niche width, and the average growth of *Streptomyces* on carbon sources it did utilize [OD>0.005] after 72 hours of growth on Biolog SF-P2 plates. Nutrient use profiles were tested with a permutational multivariate analysis of variance stratified by block niche width and the average growth were tested with a linear regression with block as a random intercept. Bold values indicate significance.

## Discussion

The ability of *Streptomyces* to utilize lignin as a sole carbon source was associated with a significant increase in the intensity of inhibition of other microbes, but only in experimental plots where crop residue was retained. Antimicrobial secondary metabolites can provide a fitness benefit to the producer by mediating resource competition but can be metabolically costly for an organism to produce and may be highly transcriptionally-regulated or lost as the cost of production increases relative to the benefit received (Kinkel et al. 2014). Our results are consistent with the hypothesis that enhanced intensity of antibiosis by lignin-utilizing *Streptomyces* results from competition-mediated selection in environments enriched with lignin-rich plant residues (Liang et al. 2023).

Extracellular lignocellulolytic enzymes can function as a public good, with their products freely available to all microbes (Allison et al. 2014). However, the extracellular oxidation of lignin is metabolically expensive, posing an energy cost to an organism that must be recuperated through the subsequent degradation of lignin-derived products and holocellulose released from lignin-bound matrices (Margida et al. 2020). We hypothesized that lignin-degrading *Streptomyces* would face selection for increased antagonistic phenotypes due to pressure to protect this costly metabolic investment. However, the lack of a significant difference in the intensity of inhibition between lignin-utilizing and non-utilizing *Streptomyces* in plots where the residue was removed suggests that selection for increased inhibition is driven by the availability of lignin-rich resources that may serve as an index to the importance of lignocellulosic resource competition to fitness.

The plant biomass-degrading microbial community is known to consist of both organisms that produce extracellular lignin-modifying enzymes (‘degraders’) and opportunistic non-lignin-degrading organisms (‘cheaters’) (Jiménez et al. 2017; Allison 2005). In communities that depend upon the degradation of complex substrates, the growth of cheaters may rely upon the diffusion of extracellular enzymes produced by degraders. In this case, degraders and cheaters may exhibit antagonism-defense arms-race dynamics, or cheaters may competitively exclude degraders, leading to the loss of both members (Allison 2005). Thus, the success of degraders faced with resource competition from cheaters relies upon the development of heterogeneous spatial structure, often brought about by biofilm formation and/or the tethering of extracellular enzymes to the degrader’s cell membrane (Chróst 1990; McKee and Inman 2019; Wakano et al. 2009; Christensen et al. 2002; Allison et al. 2014). While our culture-dependent approach captures only a subset of this community relative to high-throughput sequencing approaches, it enables the direct linkage of lignin utilization with antagonistic phenotypes at the individual isolate level. Here, the elevated intensity of inhibition by lignin-degrading *Streptomyces* under long-term residue incorporation may serve as a mechanism to defend extracellular lignin degradation from opportunistic cheaters and/or promote the development of antimicrobial metabolite-mediated heterogenous spatial structure.

Lignin-degrading *Streptomyces* have been shown to require higher levels of available nitrogen for ligninolytic activity than white rot fungi (Tuomela 2000). This may help to explain increased prominence of lignin-degrading *Streptomyces* in plots that received annual applications of nitrogen and had higher soil nitrogen levels than plots with no nitrogen amendments. The reliance of lignin-degrading *Streptomyces* on high levels of nitrogen may in part explain the greater antagonistic activities among lignin-degraders when fertilizer was not applied: competition for limited nitrogen resources could further enhance selection for antagonistic phenotypes as lignin-utilizing *Streptomyces* compete for limited nitrogen. Previous work comparing the frequency and intensity of antagonistic phenotypes among actinomycetes in long-term fertilizer amended vs non-amended plots found actinomycetes from non-amended plots had a higher frequency of inhibition (proportion of isolates which could inhibit a standard) than actinomycetes from amended plots, suggesting a connection between competition for nitrogen and the frequency of inhibition (Gieske and Kinkel 2020). However, our results show fertilization does not influence the frequency or intensity of inhibition among *Streptomyces* that cannot utilize lignin, suggesting a potential interaction between competition for nitrogen and competition for lignin. While the intensity of inhibition is not enriched among lignin-utilizing *Streptomyces* from fertilized/residue-removed plots, the proportion of *Streptomyces* that can degrade lignin is significantly enhanced relative to non-fertilized plots. This may be, in part, because the availability of nitrogen facilitates the degradation of lignocellulosic materials associated with belowground plant material, as only aboveground residue was removed.

Resource competition can facilitate antagonistic coevolution among competing microbes (Kinkel et al. 2014). While we did not test the interactions of lignin-degrading *Streptomyces* with sympatric, lignin-degrading organisms, the observed increase in the intensity of inhibition towards a single set of isolates, as well as the increase in resistance among lignin-degrading *Streptomyces* to antibiotic inhibition is consistent with a coevolutionary arms race. Indeed, the two antibiotics to which lignin-degrading *Streptomyces* exhibited significantly elevated resistance are known to be produced by genera that are known to degrade lignin and may be members of consortia that cooperate to degrade lignin (Majumdar et al. 2014). Lignin-degrading *Streptomyces* may face selective pressure to maintain resistance against antibiotics produced by commonly co-occurring microbes, even if the species may be cooperating. In contrast to lignin-degrading *Streptomyces*, the inhibition and resistance phenotypes of non-degrading *Streptomyces* are not influenced by fertilization or residue incorporation suggesting a more generalist lifestyle or the association of non-degrading *Streptomyces* with the rhizosphere (Olanrewaju and Babalola 2019).

While the ability to utilize lignin as a sole carbon source may open up a niche for *Streptomyces*, increases in constitutive antibiosis may necessitate fitness tradeoffs (D. C. Schlatter and Kinkel 2015), potentially amplified by the maintenance of lignin-modifying enzymes within the genome. However, lignin-utilizing *Streptomyces* exhibited more growth (as OD) on utilized carbon sources than non-lignin utilizing isolates while the total number of carbon sources that could be metabolized remained unchanged. Previous work suggests that the substantial energetic investment required by the oxidative degradation of lignocellulosic materials may limit the carbon use efficiency of lignin-degrading microbes, requiring lignin degrading microbes to efficiently integrate carbon into their biomass (Roller and Schmidt 2015; Moorhead et al. 2013). Indeed, lignin-utilizing *Streptomyces* exhibited more efficient growth than non-utilizing *Streptomyces* on 74 of 95 carbon sources. However, differences in the metabolic profiles of lignin-utilizing and non-utilizing isolates were primarily driven by D-gluconic acid, a naturally-occurring organic acid and a potential byproduct of the oxidation of lignocellulose (Baraniak et al. 2002). This observation suggests that the production of D-gluconic acid during lignocellulose oxidation may impose selective pressure for its efficient reuptake and metabolism, thereby minimizing carbon loss and improving carbon use efficiency.

Plant residue is a critical reservoir for the survival of overwintering plant pathogens (Bockus and Shroyer 1998). The disruption of these reservoirs, including through tillage and the burning of crop residues, can significantly reduce plant disease in subsequent crops (Bockus and Shroyer 1998; Bockus 1983). However, modern agricultural practices have sought to limit tillage to minimize erosion and improve soil health while the risks of burning include air and climate concerns, indiscriminate harm to the soil microbiome, and the risk of wildfires through uncontrolled burns (Reicosky and Saxton 2007; Holder et al. 2017; Nelson et al. 2022; Ganteaume et al. 2013). To date, efforts to harness the antagonistic potential of *Streptomyces* for the suppression of plant disease have focused predominantly on the rhizosphere and phyllosphere (Vergnes et al. 2020; Samac et al. 2003; Huang et al. 2024; Chen et al. 2018; Getha et al. 2005; Sari et al. 2021; Abbasi et al. 2021; Harsonowati et al. 2017). Here, we demonstrate that resource competition for lignin, introduced through plant litter, can contribute to selection for increased intensity of antagonism among lignin-degrading *Streptomyces*. In total, lignin-utilizing *Streptomyces* may offer multifunctional pathways for the suppression of plant disease via direct inhibition of pathogens and the potential for accelerated degradation of infectious crop residues (Perez et al. 2008). Future work should evaluate how lignin-degrading *Streptomyces* influence the persistence of infectious crop residues from senescence to pathogen germination, as well as the impact of antimicrobial metabolites on microbial survival within litter. This need is highlighted by recent work suggesting that pathogen infection can slow litter decomposition, potentially increasing the persistence of infected crop residues (Lane et al. 2026). These findings underscore the value of microbial communities, such as *Streptomyces*, that may simultaneously accelerate residue decomposition and suppress plant pathogens. Improved understanding of the interactions among fertilizer application, lignin degradation, and microbial antagonism will help optimize the use of residue-degrading, pathogen-suppressive Streptomyces as biological control agents in agronomic systems.

## Acknowledgements

We thank Julia Ahlborn, Jon Anderson, Mariah Dorner, Zoe Hansen, Aspen Hughes, Scott Klasek, Matt Michalska-Smith, and Lindsey Otto-Hanson for their research assistance.

## Data availability statement

All data generated or analyzed during this study are included in this published article (and its supplementary information files). All code is available for peer review at https://github.com/laneb14/Competition-for-lignin-and-Streptomyces-antagonism/. Upon acceptance, the code will be made available via Zenodo and a DOI will be provided.

