## Supplemental File 1 for "Incorporation of plant residue in corn monoculture selectively enriches antagonistic phenotypes among lignin-degrading vs non-degrading *Streptomyces*"

Table S1. Antibiotic discs and concentrations used for resistance assays. All "BBL Sensi-Discs" were obtained from Becton, Dickinson and Company.

| Antibiotic | Concentration (μg) |
|---|---|
| Amoxicillin / Clavulanic acid | 20 / 10 |
| Chloramphenicol | 30 |
| Erythromycin | 15 |
| Cefepime | 30 |
| Neomycin | 30 |
| Novobiocin | 30 |
| Rifampin | 5 |
| Streptomycin | 10 |
| Tetracycline | 30 |
| Vancomycin | 5 |

Table S2. Pagel's lambda statistic on the phylogenetic clustering of the 16S region of Streptomyces isolates based on fertilizer and residue practices, block from which Streptomyces were isolated, and the ability to utilize lignin as a sole carbon source. Bold values indicate significance.

| | Lambda | Log-likelihood | Likelihood ratio | P-value |
|---|---|---|---|---|
| Residue | 0.381 | -99.840 | 0.638 | 0.424 |
| Fertilizer | 0.581 | -97.627 | 5.106 | **0.0238** |
| Lignin | 0.780 | -80.689 | 19.079 | **1.254E-05** |
| Block | 7.331E-05 | -100.180 | -0.00658 | 1 |

Table S3. Post-hoc analyses comparing inverse square root intensity of inhibition (in inhibitory interactions) of *Streptomyces* which can and cannot utilize lignin. Comparisons were conducted within treatments. The block from which isolates were obtained was used as a random effect. Bold values indicate significance.

| Residue | Fertilizer | Average zone of inhibition† | F-value | P-value | Cohen's D |
|---|---|---|---|---|---|
| Retained | All Treatments | 7.43±0.639 | 35.432 | **1.26E-07** | 1.381 |
| | Applied | 5.49±0.441 | 6.961 | **0.0154** | 1.020 |
| | None | 8.55±0.934 | 29.493 | **3.09E-06** | 1.602 |
| Removed | All Treatments | 6.10±0.550 | 0.3134 | 0.578 | 0.272 |
| | Applied | 4.98±0.279 | 0.0015 | 0.969 | 0.0694 |
| | None | 7.22±1.034 | 0.3456 | 0.561 | 0.391 |

† mean±se

Table S4. Binomial generalized linear model (left) and robust linear regression (right) of the impact of fertilizer application and residue removal/retention on the proportion of standards inhibited (left) and intensity of inhibition (right). Analyses were conducted within the ability or inability to utilize lignin as a sole carbon source. Bold values indicate significance.

| | | Number of standards inhibited | | Intensity of inhibition (zone size) | |
|---|---|---|---|---|---|
| | | Z-value | P-value | T-value | P-value |
| Lignin utilizing *Streptomyces* | Fertilizer | 0 | 1 | -0.0369 | 0.970 |
| | Residue | 2.058 | **0.0396** | 6.551 | **1.14E-08** |
| | Fertilizer x Residue | -2.007 | **0.0448** | -3.016 | **0.00306** |
| Non-utilizing *Streptomyces* | Fertilizer | 0 | 1 | -0.713 | 0.514 |
| | Residue | 0 | 1 | -1.996 | 0.0720 |
| | Fertilizer x Residue | -0.487 | 0.626 | 0.941 | 0.357 |

Figure S1. Average zone of inhibition by each antibiotic disc for interactions which are not completely resistant. Bars indicate standard error. Stars indicate significance: *p<0.05, **p<0.01 ***p<0.001.

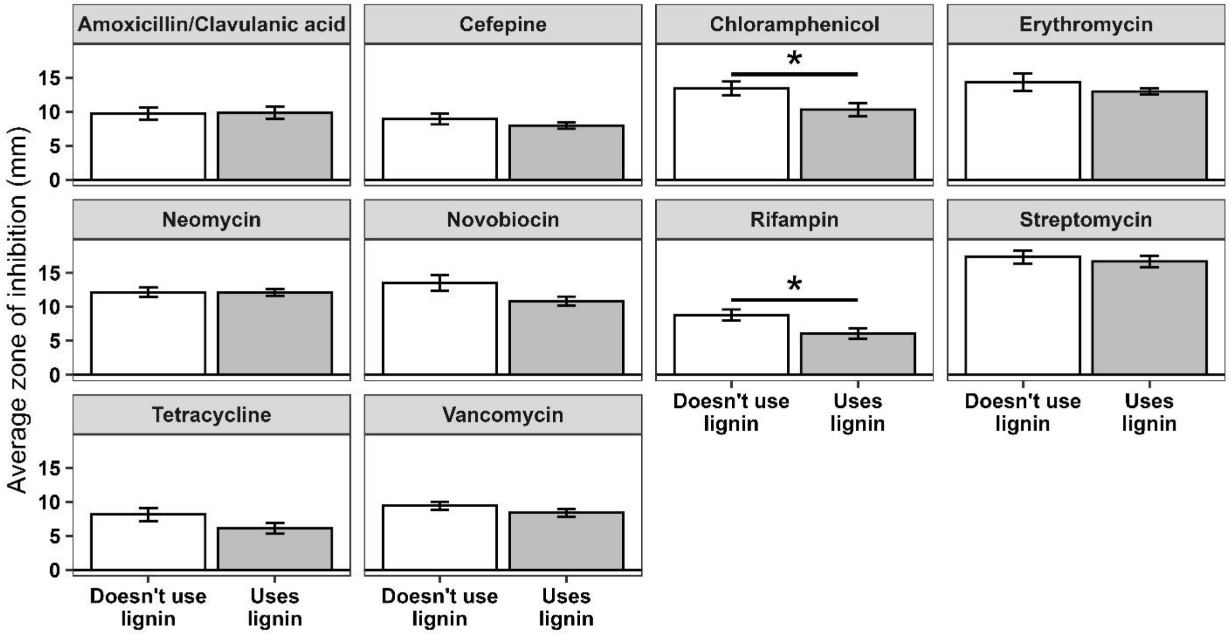

Figure S2. Violin plot of the growth of *Streptomyces* (as OD) on D-gluconic acid after 72 hours by the ability of isolates to utilize lignin as a sole carbon source. The other 94 (of 95) carbon sources did not significantly differ in growth based on the ability of isolates to utilize lignin.

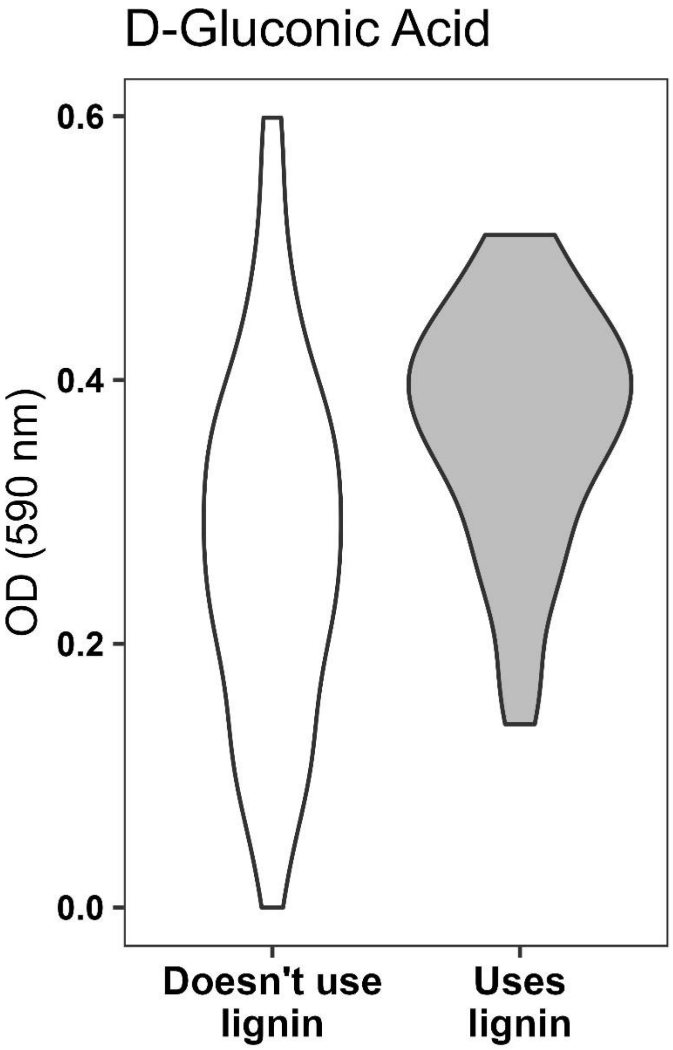
